# Unique pharmacology of mGlu homo- and heterodimers

**DOI:** 10.1101/2024.11.18.623856

**Authors:** Tyler W. McCullock, Tyler Couch, Paul J. Kammermeier

**Author notes:** Corresponding author., Department of Pharmacology and Physiology University of Rochester Medical Center, 601 Elmwood Avenue, Box 711, Rochester, NY 14642. Conflict of interest statement: Authors have no financial conflicts to declare. Although no large data sets were generated in this study, data will be made available (excel spreadsheets and Prism files) upon request.

## Abstract

**Background and Purpose:** Metabotropic glutamate receptors (mGlus) are obligate dimer G protein coupled receptors that can all homodimerize and heterodimerize in select combinations. Responses of mGlu heterodimers to selective ligands, including orthosteric agonists and allosteric modulators, are largely unknown.

**Experimental Approach:** The pharmacological properties of each group II and III mGlu homodimer (except mGlu6) and several heterodimers were examined when stochastically assembled in HEK293T cells, or specifically measured using an improved G protein mediated BRET assay employing complimented fragments of NanoLuciferase.

**Results:** Stochastically assembled receptors adopted unique signaling characteristics. Some favored the potency, efficacy or signaling kinetics of a dominant subunit, while others exhibited blended profiles reflective of a combination of homo- and heterodimers at various ratios of expressed receptor. Finally, group II and III mGlu dimers were examined for responses to selective agonists and allosteric modulators. Effects of glutamate and selective group II and III orthosteric agonists were found to result in unique concentration response profiles when examining each combination of group II and II mGlu. Effects of select allosteric modulators were examined for each mGlu2 containing dimer as well as several group III dimer pairs. Likewise, allosteric modulator effects were often unique across dimers containing the targeted subunit of the ligand being tested.

**Conclusions:** Results demonstrate that mGlu dimers respond uniquely to selective ligands, and show that the mGlu family is not governed by generalizable rules dictating consequences of dimeric subunit interactions leading to signaling consequences.

## Introduction

Metabotropic glutamate receptors (mGlus) belong to the class C of the G protein coupled receptor superfamily, and consist of 8 members (mGlu1-8), organized by sequence homology, signaling effectors, and general localization (Gregory, 2021). As class C GPCRs, mGlus are unique in a number of ways. First, they form stable, covalently linked dimers. Indeed, unlike any of the class A GPCRs, dimerization is required for mGlu function (El Moustaine *et al*, 2012). Further, while all mGlus can function as homodimers, all can also form heterodimers in some combinations, and heterodimerization can alter the pharmacological responses of mGlus in non-generalizable ways (Doumazane *et al*, 2011; Du *et al*, 2021; Goudet *et al*, 2005; Kammermeier, 2012; Levitz *et al*, 2016; McCullock & Kammermeier, 2019; Wang *et al*, 2023; Yin *et al*, 2014). Indeed, while the impact of mGlu heterodimerization is beginning to be realized, comprehensive assessment of the effects of specific ligands on mGlu heterodimers has been elusive, even as the presence of more heterodimers *in vivo* are beginning to be discovered (Meng *et al*, 2022; Moreno Delgado *et al*, 2017; Xiang *et al*, 2021; Yin *et al*., 2014).

Because of their widespread expression in the nervous system, mGlus participate in many neuronal processes and in many physiological and pathophysiological behaviors. For this reason, mGlus have been considered potential therapeutic targets for a wide range of pathologies including addiction, epilepsy, schizophrenia, and Parkinson’s disease (Ritzen *et al*, 2005). However, despite the existence of several selective and efficacious mGlu-targeting compounds entering clinical trials, they have generally not proven successful in treating human disease (Litman *et al*, 2016; Stoppel *et al*, 2021). This suggests the existence of some unknown, but possibly shared, disconnect between pre-clinical studies and successful translation to humans that may be due, at least in part, to the difficulty in predicting responses of mGlu heterodimers to these compounds. The absence of generalizable rules governing how selective compounds alter the signaling of targeted mGlu subunits across various heterodimer species can be vexing, as it makes responses difficult to predict. However, the diversity of responses to a ligand across heterodimer species could potentially reveal compounds that yield highly specific responses in vivo, where a small subset of mGlu dimers affected by a ligand intersects with restricted expression in the nervous system to give beneficial effects. Therefore, at the present time, a broad cataloguing of the effects of mGlu targeting ligands could be a fruitful approach.

Here we utilize optimized adaptations of state of the art bioluminescence resonance energy transfer (BRET) assays to study the signaling and pharmacology of mGlus when co-expressed in pairs, allowing them to be assembled stochastically as they would *in vivo*. Finally, to gain a deeper understanding of the effects of heterodimerization of mGlu pharmacology, we make specific measurements of G protein recruitment to group II and group III containing homo- and heterodimers within these populations using a technique adapted from the previously described CODA-RET approach (Urizar *et al*, 2011). Here we report for the first time isolated activity of several mGlu heterodimer species, and novel responses of many group II and III mGlus to selective orthosteric agonists and allosteric modulators. The results of these studies demonstrate that each mGlu species exhibits unique pharmacology that to date has gone unreported. These unique signals were largely unpredictable, and demonstrate the importance of making careful, dimer specific measurements to accurately assess the effects of selective orthosteric and allosteric ligands on the full complement of possible mGlus.

## Methods

### Plasmids and Molecular Biology

Plasmids encoding G_oA_, was a gift from Dr. Cesare Orlandi (University of Rochester) (Masuho *et al*, 2015b). Plasmids encoding Gβ1, Gγ2, masGRK3ct, were gifts from Dr. Stephen Ikeda (NIAAA), and the pmVenus-N1 plasmid was a gift Dr. Steven Vogel (NIAAA). The CMV-hEAAT3 plasmid was a gift from Susan Amara (NIMH; Addgene plasmid #32815). A plasmid encoding for NanoLuc was a gift from Dr. John Lueck (University of Rochester) (Ko *et al*, 2022). The plasmid encoding for the mVenus-miniGo with an N-terminal nuclear export sequence was a gift from Dr. Nevin Lambert (Augusta University). All plasmids were verified by full sanger sequencing before use.

The Gβγ-masGRK3ct sensor components, mVenus(156-239)-Gβ1, mVenus(1-155)-Gγ2, and masGRK3ct-NL were assembled to be identical to those previously reported (Hollins *et al*, 2009), with the exception of replacing RLuc8 with NanoLuc. For masGRK3ct-NL, the previously assembled masGRK3ct constructs (amino acids 495-688 of bovine GRK3 with the myristic acid sequence, MGSSKSKTSNS added to the N-terminus) and NanoLuc were copied from their original plasmids with PCR with appropriate overhangs for Gibson assembly into the EcoRV site in pCDNA3.1(+). A GCCACC Kozak sequence was added before the start codon of masGRK3ct. A GGG linker was incorporated into both overhang and both fragments. Next, pCDNA3.1(+) was digested with EcoRV, and the digested pCDNA3.1(+) was added along with the masGRK3ct and NanoLuc PCR products into an NEBuilder reaction. The reaction product was then transformed into XL10-Gold Ultracompetent E.Coli cells and colonies were screened for successful assembly. The mVenus(156-239)-Gβ1 and mVenus(1-155)-Gγ2 were cloned with an identical procedure with the incorporation of GGSGGG linker in the overhangs between the mVenus fragments and protein.

The human mGlu constructs used here were synthesized by GenScript in fragments and assembled in lab. Each hmGlu coding sequences was domesticated by eliminating all BsaI, BbsI, BsmBI, SapI, and AarI restriction sites by introducing silent mutations. Each coding sequence was then divided into two, with a GCCACC kozak sequence being added to the first fragment immediately before the start codon, the native stop codon was changed to a TGA stop, and overhangs were added for Gibson assembly into the EcoRV site of pCDNA3.1(+). Each fragment was synthesized by GenScript, and once received, the appropriate fragments were mixed with EcoRV digested pCDNA3.1(+) and subjected to a NEBuilder reaction.

To develop the NanoBit tagged hmGlu constructs, each mGlu plasmid was linearized using inverse PCR with a common forward primer that annealed to the gene’s stop codon and subsequent vector sequence and a hmGlu specific reverse primer. The reverse primer incorporated a GGSGG linker as an overhang. For LgBit tagged receptors, the LgBit sequence was then copied with PCR with overhangs corresponding to the GGSGG linker and the stop codon and vector sequence. The linearized hmGlu plasmids were then mixed with the LgBit PCR fragment and assembled with an NEBuilder reaction. For hmGlu4-LgBit, the sequence of the last 10 amino acids of the hmGlu4 c-tail were added to the LgBit reverse PCR primer to better promote its cell surface trafficking. To develop the SmBit tagged hmGlu, the hmGlu plasmids were linearized with inverse PCR using the reverse primer describe above and a modified forward primer that included the SmBit as an overhang. The resulting PCR products were then phosphorylated and ligated with an NEB KLD reaction.

To measure surface expression of the cotransfected mGlu subunits, epitope tags for HA (YPYDVPDYA) and Myc (EQKLISEEDL) were inserted after the signal peptide using Q5 site-directed mutagenesis (New England Biolabs). Tags were inserted after A22, G32, Y38, and Y36 for mGlu2, 4, 7, and 8, respectively.

After transformation of the constructs into XL-10 Gold Ultracompetent E. Coli cells and colony screening, plasmids from positive clones were isolated using an OMEGA plasmid mini prep kit and sent for Sanger sequencing to confirm successful assembly. Once sequences were confirmed by sequencing, correct clones were then transformed in subcloning efficiency DH5⍺ E. Coli cells. Transformed cultures were than propagated in 100 mL cultures and plasmids were then isolated using a Zymo midi prep kit. After isolation, the plasmids were diluted to 1 μg/μL in Tris-EDTA buffer and store in the fridge. Final plasmids were once again sent for Sanger sequencing covering the entire coding sequence to ensure no unintended mutations had previously gone unnoticed.

### HEK293T cell culture and transfection

HEK293T cell cultures were maintained in growth media consisting of Dulbecco’s Modified Eagle Media (DMEM) supplemented with 10% fetal bovine serum (FBS), 2 mM L-alanyl-L-glutamine (1x GlutaMax), 100 units/mL penicillin, and 100 μg/mL streptomycin at 37°C in 5% carbon dioxide. Cells were routinely harvested, counted, and replated every 2-3 days to prevent cultures from overgrowing. Prior to counting, cells were resuspended in Trypan Blue stain to allow for assessment of cell death, which was routinely under 10%. When conducting experiments cells were plated as described in the individual assay protocols in growth media 4 hours prior to transfection. To transfect cells, cDNA was combined polyethylenimine (PEI) in unsupplemented DMEM for 20 minutes before addition to cells. The amount of PEI added was adjusted based on the total amount of cDNA used for the transfection, using 4μL of 7.5 mM PEI per 1μg of cDNA. For some assays, the media on cells was changed to DMEM supplemented with 2% FBS only immediately prior to addition of the transfection mixture.

### NanoBRET experiments using the Gβγ-masGRK3ct sensor

For NanoBRET experiments, a modified standard protocol based on previously published protocols (Masuho *et al*, 2015a), or the mGlu optimized protocol described here were conducted. The day before the assay, HEK293T cells were plated into 6 well plates at 2 million cells per well in 1.5 mL of growth media. Four hours after plating transfections containing 200 ng masGRK3ct-NL, 200 ng mVenus(156-239)-Gβ1, 200 ng mVenus(1-155)-Gγ2, 400 ng EAAT3, 600 ng G⍺ protein, and 800 ng of receptor was assembled in 500 uL of supplemented DMEM with an appropriate amount of PEI. After 20 minutes, the transfection mixture was added to the cells dropwise. If the mGlu optimized protocol was being used, the media on the cells was changed to 1.5 mL of DMEM with 2% FBS immediately prior to adding the transfection. Cells were then allowed to transfect overnight.

The following day, the cell media was removed, and the wells were washed once with phosphate buffered saline (PBS) with no calcium or magnesium. The PBS was then aspirated and PBS with 5 mM EDTA was added. Cells were then incubated in PBS with EDTA at 37°C for 5 minutes. Cells were then harvested by titration and collected into microcentrifuge tubes. Cells were then pelleted and washed three times with imaging buffer consisting of 136 mM NaCl, 560 μM MgCl2, 4.7 mM KCl, 1 mM Na2HPO4, 1.2 mM CaCl2, 10 mM HEPES, 5.5 mM Glucose.

Experiments conducted with Low Cl-buffer used the same imaging buffer except the NaCl was replaced with 136 mM sodium gluconate. After the third wash, the cells were resuspended in appropriate volume of imaging buffer and 25 μL of cells were transferred into each well of opaque, flat bottom, white 96-well plate. For the standard procedure, cells were than assayed immediately. For the mGlu optimized protocol, cells were allowed to incubate in the plate for 1 hour before being assayed.

Cell responses were assayed using a PolarStar Omega multimodal plate reader (BMG Labtech) equipped with dual emission PMTs and two compound injectors. To select for NanoLuc and mVenus light, a 485/15 and a 535/30 filters were used respectively. Luminescent signals were integrated for 200 ms time bins, with the gain for both detectors set to 2000. Injectors were loaded with either NanoGlo reagent (1:250 dilution in imaging buffer) or test compounds and addition was automated by the plate reader. Injections were done at a speed of 430 μL per sec. For kinetic dose response experiments, 60 second time courses were used with 25 μL of NanoGlo being injected at 1 second and 50 μL of 2x test compound being added at 20 seconds. For Galpha profiling experiments, 120 second time courses were used, with injections at the same time points.

### Measuring receptor surface expression by flow cytometry

The Myc and HA tagged mGlu subunits were transfected with the other nanoBRET components as before except scaled to a 12-well plate (40% of the amounts used for a 6-well plate). The next day, cells were dissociated with FACS buffer (Dulbecco’s PBS containing 0.3% bovine serum albumin [BSA] and 5 mM EDTA) and collected by centrifugation. Cells were re-suspended in 100 µL of FACS buffer containing 5% BSA, 200 ng of Alexa Fluor 647 anti-c-Myc Antibody (BioLegend, clone 9E10), and 300 ng of PE/Cyanine7 anti-HA.11 Epitope Tag Antibody (BioLegend, clone 16B12). The amounts of antibody used were determined to have the highest staining index by serial dilution. Cells were stained on ice for 30 minutes protected from light. Cells were washed twice with 1 mL of FACS buffer, then resuspended in 300 µL of FACS buffer containing 0.2 µg/mL DAPI (Cell Signaling Technologies).

Samples were run on a FACSymphony A1 (BD Biosciences) following instrument compensation for Alexa Fluor 647 and PE/Cyanine7 collecting 20,000 events per condition. DAPI, Venus, PE/Cyanine7, and Alexa Fluor 647 were excited with a 405, 488, 561, and 633 nm lasers, respectively, and fluorescence emission was measured with 431/28, 530/30, 780/60, and 670/30 nm bandpass filters, respectively. To relate the fluorescence intensity of the PE/Cyanine7 and Alexa Fluor 647 channels, these channels were calibrated with Quantum™ Simply Cellular® (QSC) anti-mouse microspheres which have a defined antigen binding capacity (ABC) at 4 different levels.

Flow analysis was performed with FCS Express 7 (DeNovo Software). Gating for single cells was accomplished using a combination of forward scatter and side scatter height, width, and area parameters. Living cells were gated for based on their exclusion of DAPI and transfected cells were gated for based on Venus fluorescence. Median fluorescence intensities of the PE/Cyanine7 and Alexa Fluor 647 signals were converted into ABC values using a standard curve from the QSC calibration.

### BRET experiments using the CODA-RET2 system

For CODA-RET2 experiments, 4 million cells were plated into a 60 mm dishes in 3 mL of growth medium. Four hours after plating, transfections containing 400 ng of mVenus-miniGo, 600 ng of EAAT3, 2500 ng SmBit receptor, and 1500 ng LgBit receptor with an appropriate amount of PEI were mixed in 1 mL of unsupplemented DMEM. For experiments including untagged receptors, the cDNA of each receptor was adjusted, keeping a total of 4000 ng receptor cDNA. After 20 minutes, the media on the cells was replaced with 3 mL of DMEM with 2% FBS, and the transfections were added dropwise.

The following day, the cells were harvested and plated as described above and responses were assayed using the same PolarStar Omega plate reader. Due to lower light output, the NanoBit-miniGo recruitment assays were conducted as endpoint assays (except for early testing experiments). To initiate luminescent signals, 25 μL of NanoGlo reagent (1:250 in imaging buffer) was manually added to the wells, and the basal BRET ratio for each well was read serially for 3 readings with a 1 second integration. The plate was then removed from the reader and 50 μL of 2x test compound was manual added to the wells. The plate was then returned to the reader and the BRET ratio for each well was then read 5 times serially. Data acquisition was managed by the Omega Control software and the Mars Analysis software, and exported to Microsoft Excel for storage and analysis.

### Data Analysis

For BRET experiments, the BRET ratio was calculated by dividing the mVenus signals (luminescence in the 535 channel) by the NanoLuc signals (luminescence in the 485 channel):

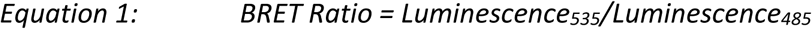

Reponses are analyzed as ΔBRET which is the average basal BRET ratio subtracted from the average stimulated BRET ratio:

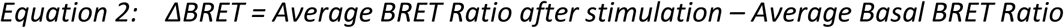

For endpoint assays, all 3 basal reading were averaged for the average basal BRET ratio, and all 5 readings post stimulation were average for the average stimulated BRET ratio. For kinetic experiments the basal BRET ratio was calculated as the average BRET ratio for 5 seconds immediately prior to test compound injection. For the average stimulated BRET ratio, the average BRET ratio for the last 10 seconds of the trace was used for non-desensitizing signals. For desensitizing signals, the average BRET ratio of a 10 second window centered at the signal’s peak was used.

### Statistics

All statistical analysis was conducted in GraphPad Prism. All ANOVA results are reported in the text in (F(dfb, dfw) = [F-value], P = [P-value]) format where dfb is the degrees of freedom between groups and dfw is the degrees of freedom within groups. Analysis reported in Figs. S2 and S3 was one-way ANOVA followed by a Holm-Šídák post-hoc test. We show the results of significance only when compared with single receptor expressed condition, as indicated in the figure legends.

## Results

### Signaling Kinetics and Pharmacology of Stochastically Assembled mGlu pairs

In a recent study, we examined the G protein coupling profiles of each of the 8 mGlu homodimers (McCullock *et al*, 2024). That study revealed differences not only in the identity of G proteins that can be activated by each mGlu, but also in the signaling properties of the various receptors. Here we aim to leverage the fast kinetics of Group II mGlus, specifically mGlu2, as a benchmark for examining responses when this receptor was co-expressed with potential dimerization partners. Figure 1A shows a schematic of the NanoBRET assay used to assess signaling properties, as well as sample responses to a range of glutamate concentrations applied to HEK293T cells expressing mGlu2, 4, 7, and 8 homodimers (and the other components of the NanoBRET system) illustrating signaling kinetics and maximal changes in BRET (Figure 1B). We previously reported that the most rapid G protein activation occurs through the group II mGlus, mGlu2&3 (McCullock *et al*., 2024). This is illustrated in Figure 1C (*upper*), which shows sample responses from indicated receptors in the NanoBRET assay to 1 mM glutamate, and in Figure 1C (lower), showing concentration response curves for each receptor. Note that 1 mM is a saturating dose in terms of both steady state activation and kinetics for each of these receptors except mGlu7, which exhibits characteristic low potency (McCullock *et al*., 2024).

**Figure 1.**
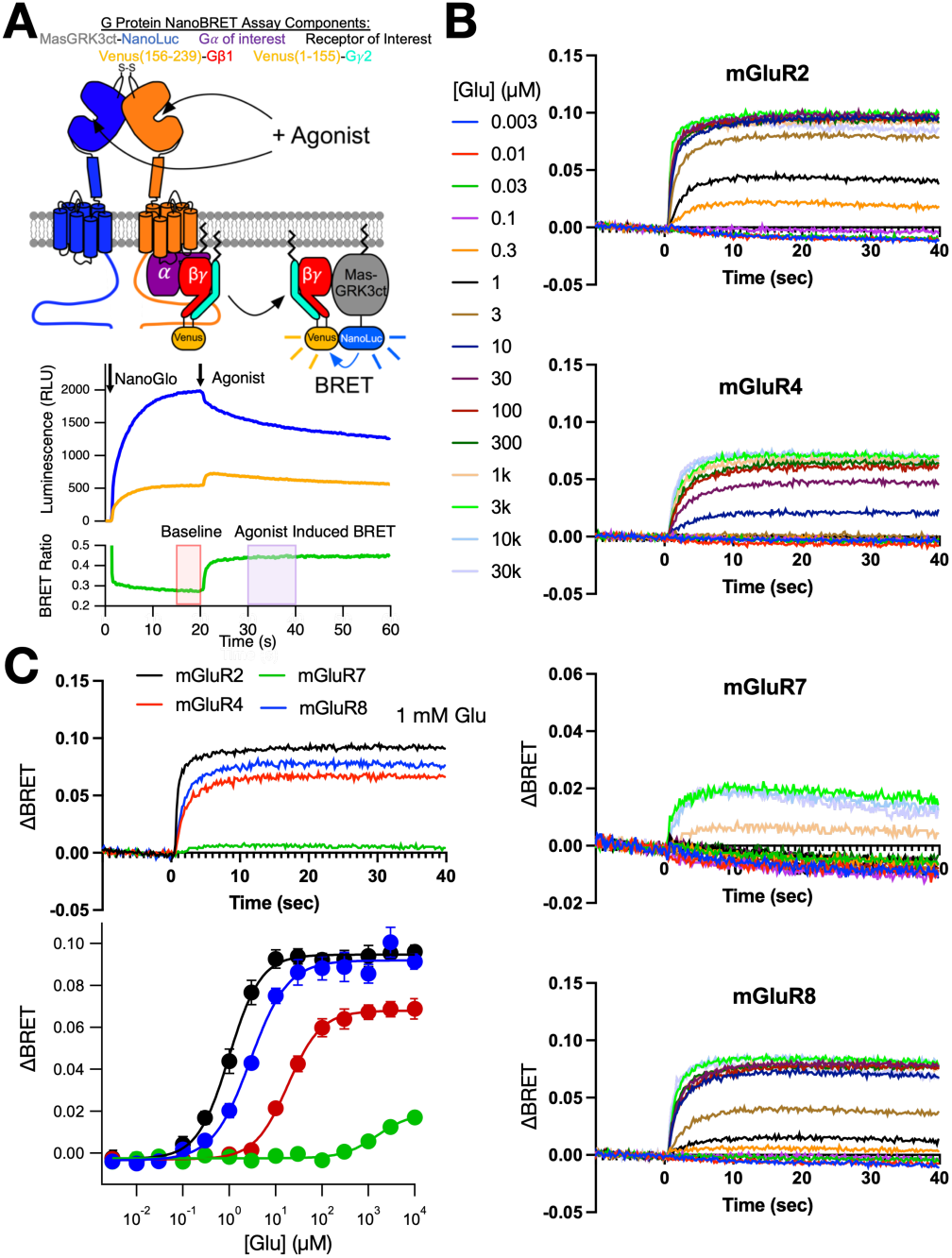
mGlu signaling measured with the NanoBRET assay. A *upper panel*, cartoon illustrating the NanoBRET assay in which receptor signaling is detected as BRET between Venus-tagged Gβγ and NanoLuc tagged MAS-GRKct, a Gβγ binding protein. *Lower panel* shows sample luminescence raw data acquired from the NanoLuc (blue) channel and the venus (yellow) channel. The ratio of these signals (V:NL) comprises the BRET measurement (green). B, Sample kinetic traces showing the change in BRET upon application of indicated [Glu] in cells expressing mGlu2, mGlu4, mGlu7, and mGlu8, as indicated. Note that 1 mM Glu represents a saturating concentration for each receptor except mGlu7. C, *upper panel* shows sample ΔBRET traces from each receptor, as indicated, illustrating that mGlu2 exhibits the most rapid activation kinetics. Lower panel shows the maximum ΔBRET from each receptor acquired as in B, as well as fits to the Hill Equation for each data set (solid lines).

These properties can be used as a guide to examine unique properties of mGlus by co-expressing several pairs of group II and III mGlus to gain a better understanding of both the propensity of specific pairs to heterodimerize as well as to begin to gain insight into the signaling properties of stochastically assembled homo-and heterodimers by expressing each pair at a number of [cDNA] ratios.

To be confident that changing the amount of cDNA for each receptor results in changes in receptor expression levels, and to assess how these changes alter receptor function, titrations of mGlu2, 4, 7, and 8 were conducted in the NanoBRET assay using G⍺_oA_ (Fig. S1) (McCullock *et al*., 2024). Concentration response curves for each cDNA concentration transfected as well as the corresponding glutamate potency and efficacy values are shown for each receptor tested. Generally, reducing the amount of cDNA of each receptor resulted in an initial reduction in potency, likely due to relief of receptor reserve effects, followed by a loss of efficacy as receptor levels presumably dropped below levels that could saturate the effector, which in this case is the MAS-GRK3-ct-NLuc detector. These results are consistent with the interpretation that reducing cDNA concentrations is an effective means to reduce receptor expression levels in this system, and can aid in the interpretation of the combined receptor results described below. We note that even when using L-AP4 for mGlu7 because of its more favorable potency, saturation of the responses could still not be achieved, so potencies for this receptor could not be accurately reported. Instead, we report basal BRET values, which reflect mGlu7’s high constitutive signaling activity (Kammermeier, 2015), as well as responses to the mGlu7 NAM MMPIP as surrogates for classical concentration response parameters to monitor mGlu7 expression. In addition to acting as a NAM on mGlu7, MMPIP exhibits inverse agonist activity that reduces the receptors constitutive signaling (Kammermeier, 2015).

We examined a series of titrations of each pair of mGlu2, 4, 7, and 8 to gain insight into the signaling and pharmacological properties of the ensembles of receptors that form when each pair is co-expressed (Figures 2, S2, and S3). Figure 2A-C shows summary data from HEK293T cells expressing mGlu2 in various ratios with either mGlu4, mGlu7, or mGlu8. We focused on mGlu2 for these studies to avoid the complication that may arise from mGlu3’s high sensitivity, and consequently high basal activation in this system (McCullock *et al*., 2024), as well as its tendency to desensitize (Abreu *et al*, 2021; Lee *et al*, 2023). Preliminary experiments suggested that mGlu2 shows higher efficacy, potency to glutamate and faster kinetics than the Group III mGlus, so expression ratios were biased away from mGlu2 to ensure that it did not obscure the signaling of the Group III mGlus. Fig. 2A (*left, center*) shows the responses to a saturating (1 mM) glutamate response in cells expressing mGlu2 and mGlu4 in 4:4, 2:6, and 1:7 ratios as well as fits to responses from each receptor in isolation (8:0 and 0:8, from Figure 1). As the amount of mGlu4 cDNA is increased, the potency of the cells glutamate responses decreases slightly, while maintaining a similar efficacy to that of mGlu2 alone (Fig. 2A). However, even at the most biased transfection conditions, the cells still showed responses more like mGlu2 than mGlu4. In contrast, the kinetics of the 1 mM glutamate stimulation clearly decreased with the addition of mGlu4 and gradually approached kinetics resembling mGlu4 (Fig. 2A *center*, quantified in Fig S2). Previous work on the mGlu2/4 heterodimer suggests that it has a shallower or slightly right shifted dose response than mGlu2 (Kammermeier, 2012; Liu *et al*, 2017). Data describing the propensity of mGlu2 and mGlu4 dimerization suggests that mGlu2/4 dimers will readily form due to mGlu4s preference for heterodimerization with mGlu2 over homodimerization, and mGlu2’s neutral preference for heterodimerization with mGlu4 over dimerization (Lee *et al*, 2020). Accounting for this, the data presented here suggest that hmGlu2 and hmGlu4 likely readily form heterodimer complexes that maintain an mGlu2 like dose response but that adopts the slower signaling kinetics of mGlu4. These results are consistent with the view that mGlu2/4 dimers signal asymmetrically by primarily coupling to G proteins through the 7TM region of the mGlu4 subunit (Liu *et al*., 2017).

**Figure 2.**
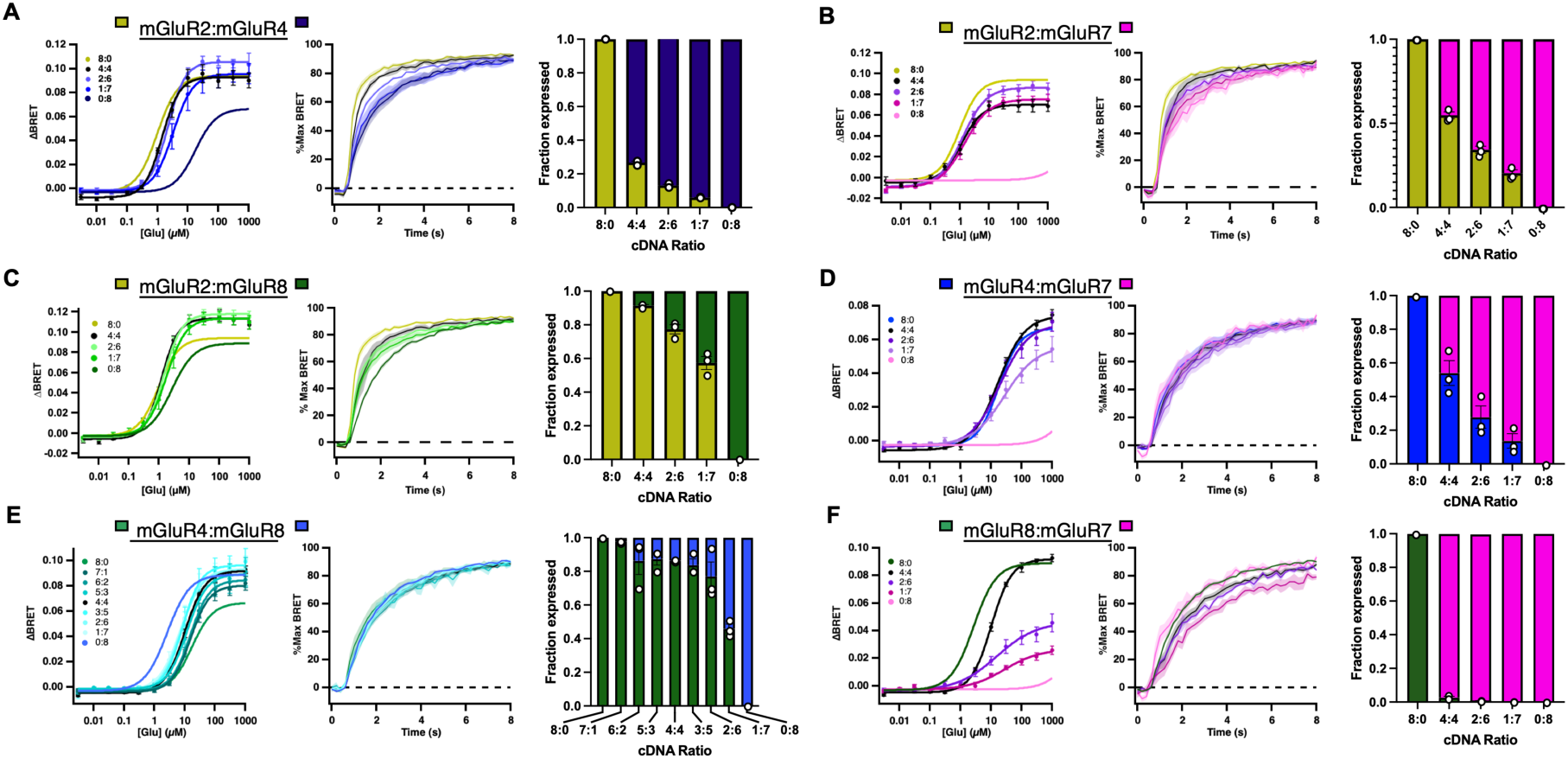
Summary of responses from stochastically assembled mGlu populations. Each panel shows glutamate dose response curves (*left*), kinetics of activation by 1 mM glutamate (*center*), and fractional expression (*right*) for a range of cDNA ratios, as indicated. A, Range of ratios of mGlu2&4. B, Range of ratios of mGlu2&7. C, Ratios of mGlu2&8. D, Ratios of mGlu4&7. E, Ratios of mGlu4&8. F, Ratios of mGlu7&8. In each panel, data from singly expressed receptors (“8:0”, “0:8”) are fits from the data illustrated in Figure 1C shown here for comparison. Overall summaries as well as results of statistical analysis are shown in Figs. S2 and S3.

To obtain a better understanding of how the observed function at each ratio reflects actual receptor surface expression levels, each receptor was N-terminally tagged (after the predicted signal sequences) with either HA or myc epitopes, and expressed at the same range of [cDNA] ratios along with all of the components of the NanoBRET system. These cells were then labeled with fluorescently tagged anti-HA and anti-myc antibodies and examined using flow cytometry. Next, the levels of each receptor were calibrated by comparing to stoichiometrically defined microspheres so that receptor surface expression levels could be directly compared (see Materials and Methods for a detailed description and Fig. S4 for gating scheme). The results of these analyses for each mGlu2 containing receptor pair are shown in Fig. 2 A-C *right*, which allows direct comparison of expression levels of each receptor to the function and pharmacology of the population. In these experiments examining mGlu2:4 ratios of 4:4, 2:6, and 1:7, the fraction of mGlu2 was 0.27±0.01, 0.12±0.01, and 0.06±0.001, respectively. A number of interesting findings are apparent. For example, when mGlu2 and 4 are expressed, even at ratios where mGlu4 expression is clearly dominant, the pharmacology of the population more closely resembles mGlu2, lending further support for the idea that mGlu4 has a strong propensity to heterodimerize with mGlu2 (Lee *et al*., 2020), as mentioned above.

Repeating these experiments with mGlu2 and mGlu7 reveals similar results (Fig. 2B). As expression was shifted towards mGlu7 a dose response similar to that of mGlu2 alone is maintained, but the kinetics of the response remain more like mGlu7 (Fig. 2B right, quantified in Figure S2). Differences in potency, efficacy, and kinetics were observed across the range of conditions (see Fig. 2B, legend). The efficacies of the responses were more variable but were significantly different from mGlu2 and mGlu7 homodimers at some ratios (Fig. S2). Analysis of expression levels using flow further highlight these striking results, as expression levels at most of the ratios tested favor mGlu7 over 2 (Fig. 2B, *right*), showing the fraction of mGlu2 when expressed at 4:4, 2:6, and 1:7 being 0.55±0.02, 0.34±0.02, and 0.20±0.02, respectively. Work describing the mGlu2/7 heterodimer suggested that it should have a glutamate dose response slightly left-shifted compared to mGlu2, based on FRET measurements of conformational changes of the extracellular domains (Habrian *et al*, 2019). This result was not replicated when co-expressing mGlu2 and mGlu7 together on a global level in this study, however. This may due the reported disfavoring of heterodimerization between mGlu2 and mGlu7, resulting in only a limited population of mGlu2/7 dimers forming upon co-expression (Lee *et al*., 2020). In those studies, it appears this population forms at such low levels, they are not able to influence the global glutamate dose response. Instead, the overt phenotype simply seems to be the suppression of mGlu2 expression in favor of mGlu7 expression, resulting in corresponding right shifts in potency, and decrease in efficacy and kinetics.

Next, responses from the co-expression of mGlu2 and mGlu8 were investigated (Fig. 2C). In general, mGlu8 is the most understudied mGlu, and to our knowledge, the only data regarding mGlu8 heterodimerization is indirect dimerization propensity data with mGlus4, 2, and 3 (Lee *et al*., 2023; Lee *et al*., 2020) and a more recent study examining potential mGlu7/8 heterodimers in the hippocampus (Lin *et al*, 2022). Interestingly glutamate dose responses from cells expressing mGlu2 and mGlu8 in 4:4, 2:6, and 1:7 ratios maintained a higher potency than mGlu8 homodimers (Fig. 2C, quantified in Fig S2). Under these conditions, the fraction of mGlu2 expressed was 0.91±0.01, 0.77±0.03, and 0.57±0.04, respectively. In addition, the kinetics of the 1 mM glutamate response maintained an intermediate phenotype for all combination conditions that did not reflect the kinetics of either mGlu2 or mGlu8 homodimers (Fig. 2C).

Further studies on isolated mGlu2/8 heterodimers may shed further light on the phenotypes observed here. Interestingly, unlike the other combinations described above, expression of mGlu2 seemed to be favored over mGlu8, which is most evident by examining the 4:4 [cDNA] ratio in each panel.

We next examined responses of the group III mGlus when expressed in combination. Although mGlu4, mGlu7, and mGlu8 exhibit similar kinetics, each has well resolved differences in their glutamate dose response parameters that should enable comparisons within the Group III family. This is a particularly attractive endeavor as there is very limited pharmacology reported for any group III heterodimer combination, probably due to the lack of subtype specific pharmacological tools. For these studies, the low potency of mGlu7 is a benchmark, allowing the biasing of transfections towards mGlu7 should allow for the isolation of signals generated by only the other receptor species as long as glutamate concentrations under 1 mM are used. In the case of mGlu4 and mGlu8, co-expression using an increased number of conditions was evaluated to make up for the lack of selective activation by glutamate and similar signaling kinetics.

Co-expression of mGlu4 and mGlu7 in 4:4, 2:6 and 1:7 ratios produced responses and kinetics similar to mGlu4 homodimers (Fig. 2D; Fig S3), even though the expression levels of both receptors were comparable at the 4:4 ratio, and mGlu7 was dominant at most ratios examined (Fig. 2D, *right*). In these experiments, the fraction of mGlu4 expressed was 0.53±0.08, 0.28±0.07, and 0.14±0.04, at 4:4, 2:6, and 1:7, respectively. Across the entire data set, significant differences were observed in efficacy, potency, and kinetics, but only comparing each condition to mGlu7 homodimers (Fig. 2D). It should be noted that these statistical tests are largely detecting the poor performance of mGlu7 compared to the remainder of the conditions. In preliminary experiments, mGlu4 responses were greatly influenced by the amount of cDNA included in the transfection, with larger amounts of cDNA producing higher glutamate potency values. This phenomenon did not plateau even when the maximum amount of mGlu4 cDNA was transfected. In addition, dimerization propensity data suggest mGlu4 and mGlu7 have a neutral preference for heterodimerization over homodimerization (Lee *et al*., 2020) which supports the formation of a significant population of heterodimers when co-expressed. Taking these observations into account, this data suggest that mGlu4/7 heterodimers may possess indistinguishable pharmacology from mGlu4.

To gain insight into the responses of mGlu4 and mGlu8 expressing cells, a wide array of mGlu4:mGlu8 expression conditions were assayed to make up for the lack of unique signaling properties between these receptors (Fig. 2E). As seen in Fig. 2E *right*, expression levels at most ratios favored mGlu4. No differences in signaling kinetics, and only minor differences in efficacy were observed across all expression ratios (Fig. 2E, quantified in Fig S3) but the dose response potencies gradually changed from an mGlu4 like phenotype towards (but not reaching) an mGlu8 like phenotype. Notably, the glutamate potencies at most ratios more closely resembled mGlu4 homodimers, even when the cDNA was skewed towards mGlu8, which probably only reflects protein expression levels. Fractional expression of mGlu4 in the ratios listed inf Fig. 2E were 0.98±0.002, 0.86±0.08, 0.87±0.04, 0.86±0.003, 0.84±0.04, 0.77±0.09, and 0.46±0.03, respectively. Work from others has suggested that mGlu4 prefers heterodimerization with mGlu8, but mGlu8 is neutral towards dimerizing with mGlu4 (and favors mGlu2 and 3) (Hofmann *et al*, 2021; Lee *et al*., 2023). These findings suggest that mGlu8 is functioning from complexes that possess either an mGlu4 like response, or an intermediate response when mGlu4 is present.

Finally, unlike mGlu4, when mGlu8 was expressed with mGlu7, the glutamate dose response changes noticeably, and appears to reflect an intermediate phenotype (Fig. 2F). This is to some degree unsurprising when looking at the expression levels (Fig. 2F, *right*). Still it is quite remarkable that at ratios where the mGlu8 expression is very low but still detectable, the pharmacology of the population remains dramatically altered from mGlu7 homodimers. These data may suggest a strong tendency of mGlu7 to heterodimerize with mGlu8. The fractional expression of mGlu8 in the 4:4, 2:6, and 1:7 expression conditions was 0.025±0.007, 0.008±0.002, and 0.003±0.001, respectively (Fig. 2F, *right*). The potency of the responses gradually decreases as the mGlu8 fraction is increased (Fig. 2F, *left*), and the efficacy of the response decreases in parallel. This may at least in part reflect a strong decrease in mGlu8 expression rather than signaling from a heterodimer, as the mGlu8 levels decrease dramatically (Fig. 2F, *right*), but the kinetics do appear to slow to become more like mGlu7 (quantified in Fig S3). Note that the difference in kinetics between mGlu7 and mGlu8 is small to begin with. In addition, it may not be completely appropriate to compare the kinetics between conditions for this set of experiments as the dose responses suggest that 1 mM glutamate may no longer be a saturating stimulus for the 6:2 and 1:7 conditions. When comparing individual conditions, the combined conditions gradually shift from being similar to mGlu8 to being similar to mGlu7, with multiple combinations being significantly different from each homodimer. Due to the lack of studies focused on mGlu7 and mGlu8 dimerization (but see (Lin *et al*., 2022)), it’s difficult to pin down the mechanism of these changes. Given the experimental set up, one of two possibilities seem most likely: either the mGlu7/8 heterodimer adopts mGlu7 like responses to glutamate, or the expression of mGlu7 suppresses mGlu8 expression, without significant formation of a population of mGlu7/8 heterodimers to support signaling. Since it appears that mGlu8 does not strongly favor dimerizing with mGlu7 (Lee *et al*., 2023), our interpretation favors the latter.

### Examining heterodimer specific signaling with CODA-RET2 reveals unique pharmacology

We next aimed to examine responses to mGlu heterodimers directly by building off the principles the Javitch lab developed with their CODA-RET (COmplemented Donor-Acceptor Resonance Energy Transfer) system to investigate the activity of dimeric receptor species specifically (Urizar *et al*., 2011). Given that CODA-RET is a BRET assay that would be amenable to high throughput technologies, we adapted this technology in an attempt to optimize performance for mGlu assays. Briefly, we replaced the split Renilla luciferase used in that study with NanoLuc, and replaced the full length, FP-tagged G⍺ protein with a venus tagged miniG⍺ (mVenus-mG⍺o) (Wan *et al*, 2018) to help enhance the BRET signal. As a validation of the system, a number of control experiments were performed with LgBit and SmBit tagged mGlu2 and mu opioid receptor (MOR: Fig S5), which should not dimerize. These experiments demonstrated that significant luminescence could only be observed accompanying the stable dimerization of mGlu2 (mGlu2-LgBit + mGlu2-SmBit), and that the BRET signal upon agonist activation was restricted to that from the agonist activating receptor with the full length or assembled NanoLuc tag (Fig. S5), ruling out crowding or bystander BRET effects.

Responses to specific mGlu ligands were examined on group II and III heterodimers using selective orthosteric agonists. To screen the pharmacology of Group II and Group III heterodimers, LgBit and SmBit tagged mGlu2, mGlu3, mGlu4, mGlu7, and mGlu8 were transfected in all combinations (25 in total) along with mVenus-mG⍺o (See Fig. 3A & B).

**Figure 3.**
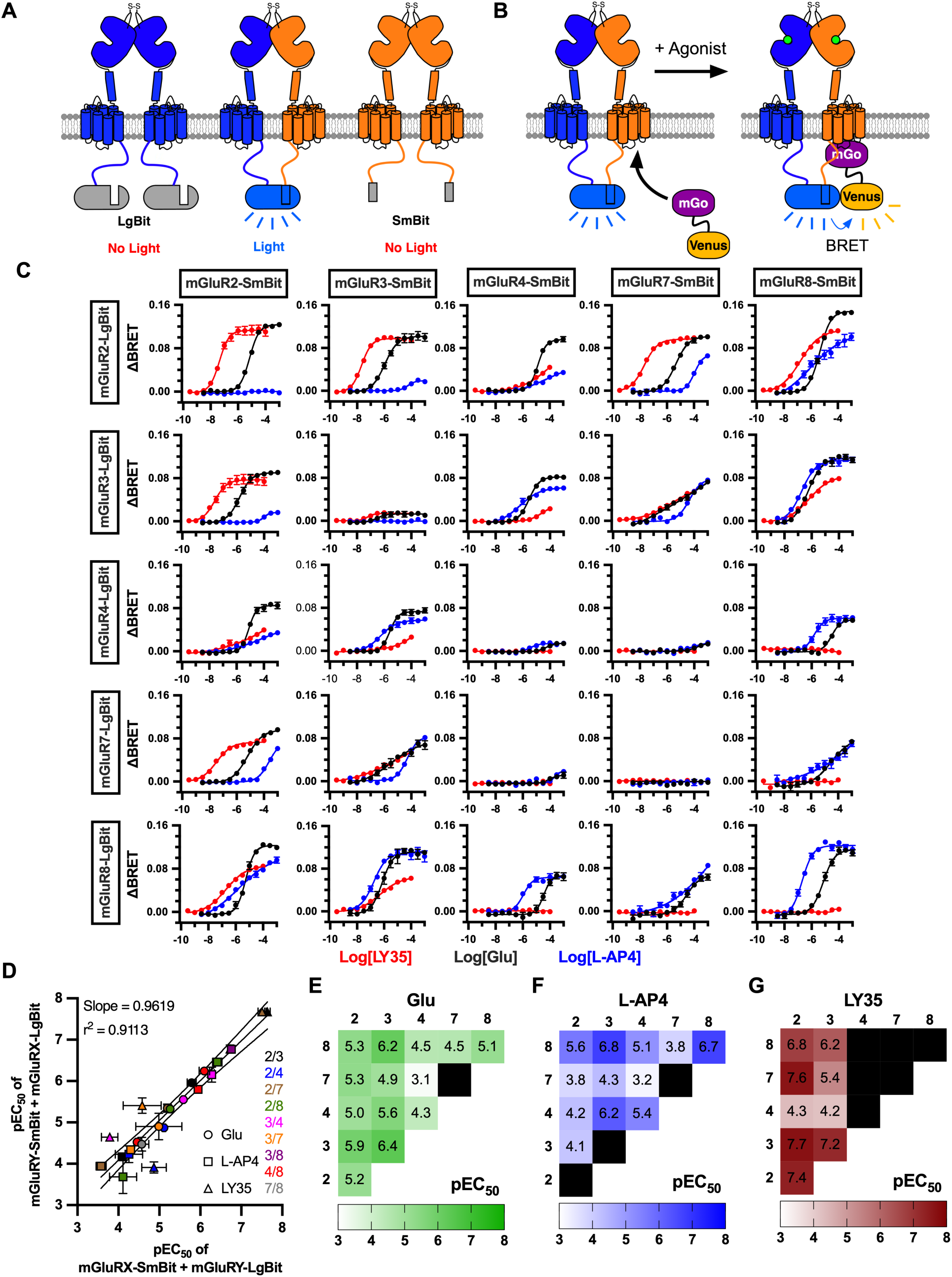
Summary of agonist responses in the CODA-RET2 assay. A, Cartoon depicting the selective generation of light of only LgBit tagged and SmBit tagged mGlu heterodimers. B, Cartoon depicting the translocation of mVenus-mG⍺o to a LgBit/SmBit mGlu heterodimer upon activation and the resulting BRET that detects this interaction. C, Reponses of all 25 combinations of Group II and Group III NanoBit mGlu receptors to glutamate (black), LY354740 (LY35, red), and L-AP4 (blue). LgBit receptors are listed to the left and SmBit receptors for each condition are indicated across the top. The Y-axis is ΔBRET, and the X-axis is Log10 [Compound (M)]. All data points are displayed as the average ± SEM but in some cases the error bars are smaller than the marker. N = 3-5 biologic replicates per expression condition per compound. D, Correlation of pEC_50_ values for each response curve for the data in C, from which a reasonable fit could be generated for each individual response. All data points display the average ± the SEM, but some error bars are smaller than the marker. The linear regression line is shown as a solid black line with the 95% confidence interval shown as dashed black lines. E-G, EC_50_ values are shown as heat maps for all combinations of mGlu homo- and heterodimers to glutamate (E), L-AP4 (F), and LY35 (G) as measured in the CODA-RET2 assay.

Experiments were performed in replicate with three different compounds: glutamate, the Group II specific orthosteric agonist LY354740 (LY35), and the Group III specific orthosteric agonist L-AP4. The use of subtype specific compounds is a keystone of studies that attempt to identify heterodimers in more complex systems (Moreno Delgado *et al*., 2017; Xiang *et al*., 2021; Yin *et al*., 2014), so developing an understanding of their behavior on each heterodimer is crucial. The resulting 75 average dose responses are displayed in Fig. 3.

Examination of the CODA-RET2 dose response data reveals some very surprising, unique pharmacological profiles of even similar heterodimers. For example, when mGlu2 dimerizes with the group III mGlus (4, 7, and 8), the rank order of potency to glutamate, L-AP4, and LY35 is different in each case (Fig. 3C, *top*). Also surprising was that when mGlu3 dimerized with each of these group III mGlus, the pharmacology of the mGlu3/x dimers was quite different than that of the mGlu2/x dimers, despite mGlus 2&3 being quite similar pharmacologically. Orthosteric agonist pharmacology also differed among the group III heterodimers. Perhaps most notably was a dramatic flattening of the curves with some of the dimers, mGlu7 containing heterodimers in particular, including mGlu7/3 but notably not mGlu7/2 (Fig. 3C).

In general, the screen showed excellent dynamic range and signal to noise ratio for most combinations of receptors. Interestingly, mGlu3 and mGlu4 both produced comparatively small amplitude changes in this assay in homomeric arrangements, but both showed considerably larger responses when in combination with other receptors, or each other.

Despite their low amplitudes, mGlu3 and mGlu4 homodimers still produced the predicted pharmacology with glutamate and the subtype specific agonists as reported elsewhere (Gregory & Goudet, 2021). This demonstrates how the high signal-to-noise ratio of this system can enable detection of responses even under non-ideal conditions. While mGlu3 and mGlu4 produced smaller signals than desired, the only true failure of the system seems to be with mGlu7 in homodimer configuration, which showed no responses in this assay to any drug but was able to generate large responses with other receptors including mGlu3, and detectible changes in BRET when with mGlu4. Note also that the mGlu7-LgBit and -SmBit constructs performed indistinguishably from wild type mGlu7 in the NanoBRET assay using L-AP4 as an agonist (not shown), suggesting that the lack of signal in the CODA-RET2 assay with mGlu7 homodimers is likely the result of a smaller signal to noise compared with the NanoBRET system, but that expression of the tagged receptors is not likely impaired.

The results shown in Fig. 3C represent duplicates of each heterodimer condition, which should be detecting the pharmacology of the same species. For example, both the mGlu2-SmBit + mGlu7-LgBit condition and the mGlu2-LgBit + mGlu7-SmBit conditions will report the responses of the mGlu2/7 heterodimer. If the CODA-RET2 system is selecting for the heterodimer responses properly, these reciprocal conditions should report identical pharmacology. To assess if either configuration of the NanoBit tags (mGluX-SmBit + mGluY-LgBit and mGluX-LgBit + mGluY-SmBit) report the same pharmacology, the EC_50_ for each drug was plotted against that in its reciprocal expression condition (Fig. 3D). Correlating the potencies of all responses that could be reasonably fit by the reciprocal conditions indicates that each of the two conditions report similar responses (Fig. 3D). Linear regression of the correlated potencies shows a slope close to 1 (0.9619) and excellent agreement between the fitted line and the data (r^2^ = 0.9113). The conditions that deviate most from the fit line are primarily lower potency responses that do not show good saturation, so it’s likely the error is in the EC_50_ estimate rather than the behavior of the receptors. Finally, heat maps and values of the potencies for Glutamate (Fig. 3E), L-AP4 (Fig. 3F), and LY35 (Fig. 3G) are also shown.

Black squares indicate no responses to the drug for the indicated receptor dimer. Summaries of the pharmacological data from all of the experiments described in Fig. 3 are shown Fig. 4. Fig. S6 shows summaries for each agonist of the E_max_ in ΔBRET (Fig. S6A), the pEC_50_ values (Fig. S6B), and Hill slopes (Fig. S6C). These results bolster the conclusion that some rather dramatic differences in pharmacological properties across ostensibly similar mGlu dimers can be observed in ways that may not be predictable by the known pharmacology of dimer subunits alone. For example, comparing responses to mGlu2/4 to 3/4 and 2/7 to 3/7 reveals stark differences despite the pharmacological properties of mGlu2 and mGlu3 homodimers being similar.

**Figure 4.**
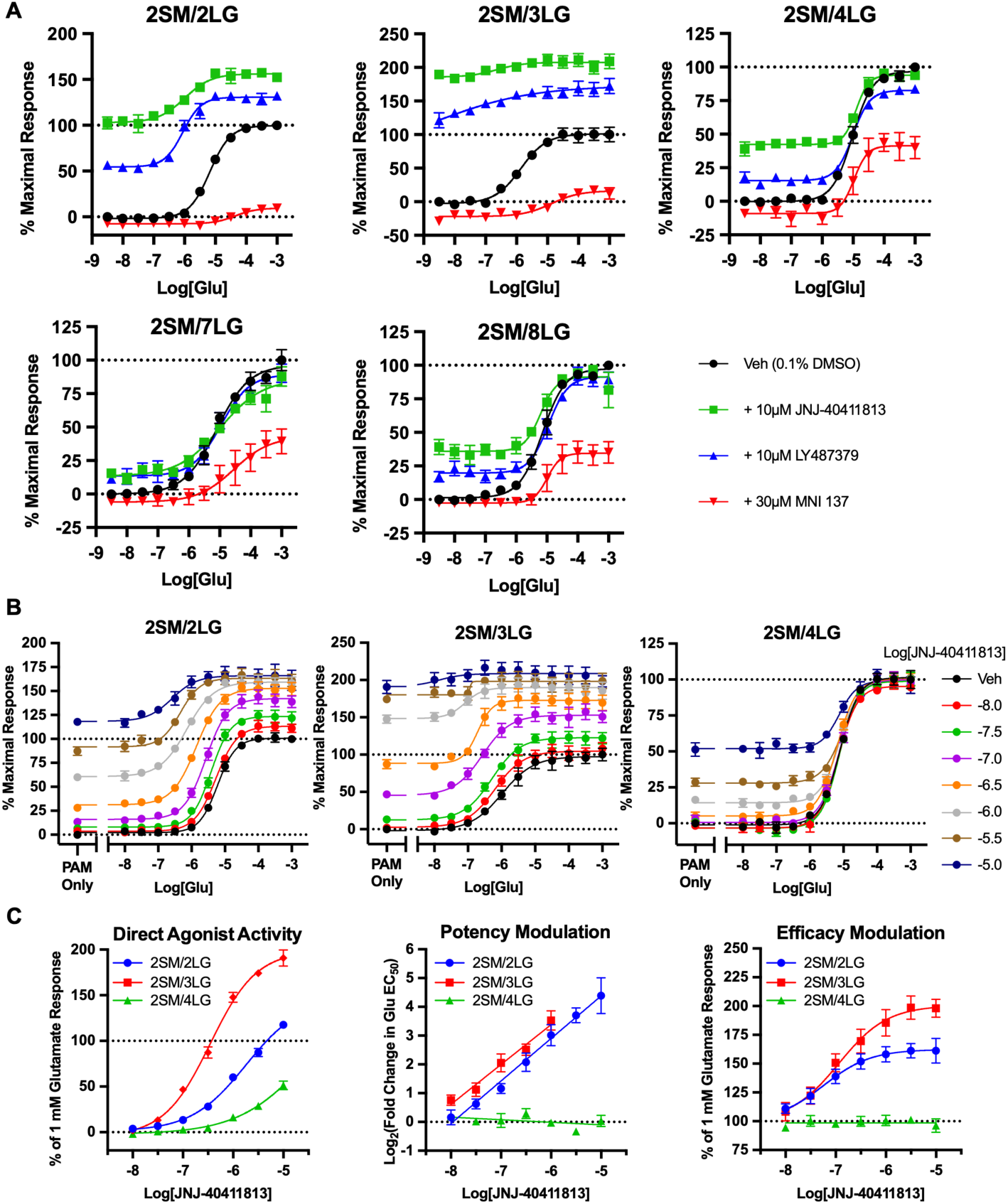
mGlu2 containing dimers respond differently to mGlu2-targeting allosteric modulators. A, Glutamate dose response curves in the presence of vehicle (black), 10 µM JNJ404 (green), 10 µM LY48 (blue), or 30 µM MNI-137 (red) acquired using the CODA-RET2 system for mGlu2 homodimers and heterodimers with mGlu3, 4, 7, and 8, as indicated. Panel B shows glutamate dose response curves for mGlu2/2, 2/3, and 2/4 in the absence (black) and presence (as indicated) of 7 concentrations of JNJ404 ranging from 10 nM to 10 µM. Summaries of JNJ404 direct agonist activity, potency modulation, and efficacy modulation are also shown in Panel C for mGlu2/2 (blue), 2/3 (red), and 2/4 (green). Note that in the center panel, data are presented in a log/log plot to emphasize differences between the mGlu2/2 and 2/3 conditions. All data points are displayed as the average ± SEM but in some cases the error bars are smaller than the marker. N = 3-5 biologic replicates per expression condition per compound.

### Specific effects of allosteric modulators on individual mGlu2 containing heterodimers

To directly examine responses to mGlu2-targeting allosteric modulators, the CODA-RET2 system was employed on mGlu2-SmBit co-expressed with LgBit tagged mGlu2, 3, 4, 7, and 8 (Fig. 4). We examined responses to two PAMs: LY487379 (LY48) and JNJ-40411813 (JNJ404), and one NAM: MNI-137. LY48 and MNI-137 are widely used AMs (PAM, and NAM, respectively) that target mGlu2 at known sites (Cid *et al*, 2010; Lundstrom *et al*, 2016; O’Brien *et al*, 2018), and JNJ404 (also called ADX-71149)(Lavreysen *et al*, 2015) is a newer mGlu2 targeting PAM that has previously been in clinical trials for the treatment of schizophrenia, and more recently for epilepsy (Cid *et al*., 2010; Kent *et al*, 2016; Metcalf *et al*, 2017). Fig. 4 shows glutamate responses with vehicle (black), and in the presence of 10 µM JNJ404 (green), 10 µM LY48 (blue), and 30 µM MNI-137 (red) for each mGlu2 containing dimer combination, as indicated.

Summaries of the glutamate potencies and efficacies in the absence and presence of each modulator are shown in Fig. S7A. The NAM MNI-137 inhibited each dimer, although more strongly on the exclusively group II dimers (mGlu2/2 and 2/3). Responses to the PAMs LY48 and JNJ404 also varied across mGlu2 dimers. The exclusively group II dimers exhibited strong direct agonist effects at this concentration of both PAMs, and strongly potentiated glutamate responses. Interestingly, direct agonist effects of both PAMs were considerably stronger on mGlu2/3 than on mGlu2/2. None of the group III containing dimers exhibited potentiation of glutamate responses with either PAM. mGlu2/4 and mGlu2/8 experienced some direct agonism by both PAMs, but this was much weaker than on mGlu2/2 or mGlu2/3. Finally, mGlu2/7 was only marginally altered, if at all, by either PAM. These data provide strong evidence that mGlu2-targeting allosteric modulators, particularly PAMs, produce distinct effects on different mGlu2 containing dimers, and in fact do not act as PAMs at all on mGlu2 dimers containing a group III subunit, but instead act only as allosteric agonists. The potency and perhaps strength of this agonism varied by subunit composition and by ligand (see below).

To further probe these distinct pharmacological profiles, responses of three dimers, mGlu2/2, 2/3, and 2/4 were examined in more detail using several JNJ404 concentrations from 100 nM to 100 µM, against a range of glutamate concentrations (Fig. 4B&C). These data illustrate the striking differences in responses of these three mGlu2 containing dimers to the PAM JNJ404.

The basic observations from Fig. 4B are clear: JNJ404 acts as a direct agonist on each dimer, potentiates glutamate responses only on mGlu2/2 and 2/3. Moreover, the strength of direct agonist activity varies dramatically across each dimer, ranging from mGlu2/3 > mGlu2/2 > mGlu2/4. The data in Fig. 4C also make it clear that saturation of direct agonist effects was not achieved for mGlu2/2 or mGlu2/4, even at the near solubility limited 10 µM JNJ404, so it remains unclear if the differences lie in potency, efficacy, or both. Interestingly however, the effects of JNJ404 on glutamate efficacy do appear to saturate (Fig. 4C). Finally, note that the graph illustrating changes on dimer potency with JNJ404 (Fig. 4C, *center*) are plotted on a log-log scale to better illustrate the differences between mGlu2/2 and 2/3. While effects of [JNJ] differ somewhat between these dimers, the offset may reflect the difference in glutamate potencies of the respective receptor subunits.

### Specific effects of allosteric modulators on individual group III heterodimers

To assess differential effects on group III heterodimers, we first examined responses to the mGlu4-targeting PAMs VU0155041 (VU015) (Rovira *et al*, 2015), VU0361737 (VU036), which bind mGlu4 at separate sites and have differential effects on the mGlu2/4 heterodimer (Engers *et al*, 2009; Fulton *et al*, 2019; Rovira *et al*., 2015), and Foliglurax (Charvin *et al*, 2017), which has recently failed in phase 2 clinical trials to reduce adverse effects of levodopa (Carvalho, 2020) (Fig. 5A). As with the mGlu2-targeting compounds, clear differences were seen using PAMs targeting mGlu4. In some cases, some differences were also seen with different PAMs on the same dimers. For example, while all the PAMs strongly potentiated glutamate responses on mGlu4/4, almost no effect was seen on mGlu4/2 and 4/3 with Foliglurax and VU036, with modest enhancement by VU015 (Fig. 5A). Similar but less pronounced effects were seen with each PAM on the group III heterodimers dimers mGlu4/7 and 4/8 compared with the mGlu4 homodimer. The effects of Foliglurax were examined in more detail by applying it to mGlu4/4, 4/7, and 4/8 in the CODA-RET2 system. Fig. 5B shows glutamate concentration response curves in the absence (vehicle) and presence of a range of Foliglurax concentrations.

**Figure 5.**
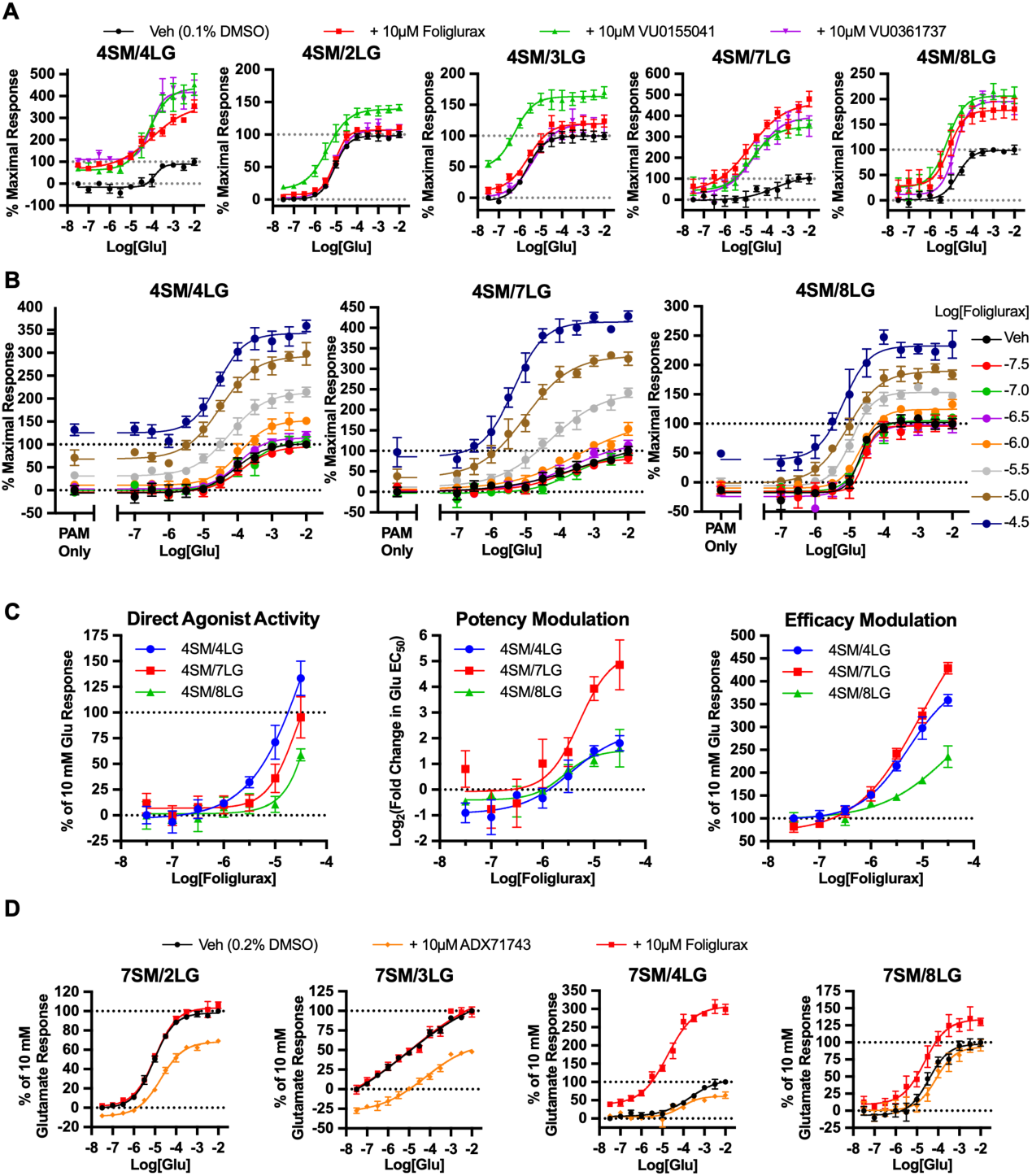
Group III mGlu dimers display unique responses to allosteric modulators targeting mGlu4 and mGlu7. Panel A shows Glutamate dose response curves in the presence of vehicle (black), or 10 µM Foliglurax (red), 10 µM VU0155041 (green), or 10 µM VU0361737 (purple) applied to the indicated mGlu4 containing dimers using the CODA-RET2 system. Panel B shows glutamate dose response curves generated in the presence of a range of [Foliglurax], as indicated for mGlu4/4 (left), 4/7 (center), and 4/8 (right). Summaries of the Foliglurax effects on these three dimers are also shown, including direct agonist effects (left), modulation of glutamate potency (center), and efficacy (right). Panel D shows Glutamate dose response curves in the presence of vehicle (black), 10µM ADX71743 (gold), or 10 µM Foliglurax (red) for the indicated mGlu7 containing dimers All data points are displayed as the average ± SEM but in some cases the error bars are smaller than the marker. N = 3-5 biologic replicates per expression condition per compound.

Summaries of the direct agonist effects and Foliglurax mediated changes in glutamate responses (Fig. 5B&C) are also shown. While Foliglurax acts as a direct agonist and potentiates the effects of glutamate on each dimer, there are differences in its effectiveness (Fig. 5B&C). Most notably, direct agonist effects on mGlu4/8 are particularly weak and require greater than 10 µM Foliglurax.

Finally, we examined effects of Foliglurax as well as an mGlu7 targeting NAM, ADX71743 at 10 µM (Fig. 5D) on several mGlu7 containing dimers: mGlluR7/2, 7/3, 7/4, and 7/8. Because of the poor responses of mGlu7/7 in the CODA-RET2 system, we did not examine effects on homodimers. Nevertheless, interesting dimer specific effects were seen with each drug. Unsurprisingly (Fig. 5A, above), Foliglurax strongly potentiated signaling of mGlu7/4, and did not alter responses to mGlu7/2 or 7/3, which without an mGlu4 subunit was expected.

However, mGlu7/8 was weakly potentiated by Foliglurax, which may suggest Foliglurax can bind mGlu8 as well as mGlu4, although this was not directly tested. ADX71743 inhibited responses of mGlu7/2, 7/3, and more weakly, 7/4, but did not detectably alter mGlu7/8 signaling. Summaries of the EC50 and Emax values for modulators targeting mGluR4 and 7 are shown in Fig. S7B&C, respectively. Our results with mGlu7/8 are in contrast to results using ADX71743 in recently published work using similar methods (Lin *et al*., 2022). The source of this discrepancy is unclear, but we would note that in our experiments, we used glutamate as an agonist while Lin et al. used DL-AP4. In addition, we note that while we used L-AP4 instead of DL-AP4, the responses we saw from mGlu7/8 (see Fig. 3) were dramatically different from what they reported. This may be an indication that there are differences in the BRET systems beyond drug effects that could account for these differences. It is somewhat surprising that ADX71743 more strongly inhibited signaling of mGlu7 when dimerized with group II mGlus than when paired with other group III receptors, which may be an indication of a different functional relationship of mGlu7 subunits when paired with the group II and group III mGlus.

## Discussion and Conclusions

### Stochastically assembled mGlus

We assessed mGlu dimer signaling using two approaches: stochastic assembly or paired receptors by co-expression of group II and III mGlus, and using the CODA-RET2 approach to examine G protein recruitment to mGlu dimers tagged with NanoLuc compliments LgBit and SmBit. mGlus have been recognized to form covalently linked heterodimers for over 10 years (Doumazane *et al*., 2011), but detailed information about these heterodimers is limited. This is largely due to the stochastic assembly of dimers when mGlus are expressed together, resulting in formation of both heterodimers and homodimers at unknown proportions, which makes Isolating heterodimer signals challenging. Several labs have developed creative solutions to this problem, but wide scale implementation of any of these solutions has yet to be realized (Habrian *et al*., 2019; Kammermeier, 2012; Levitz *et al*., 2016; Werthmann *et al*, 2020; Xiang *et al*., 2021). Compounding this issue, these studies used different technologies to investigate these heterodimers making it difficult to compare data across studies. Another important consideration is whether a reductionist approach is always most appropriate. Studies measuring mGlu heterodimer pharmacology in isolation will improve our understanding, but physiologically stochastic assembly is likely to occur *in vivo*. Certain dimer species may accumulate at particular synapses (Xiang *et al*., 2021), but this evidence is limited and more direct support will be needed to confirm these observations.

Therefore, responses of cells expressing mGlus in different ratios were examined here. The unique signaling properties described in Fig. 1 (efficacy, potency, and kinetics) served as a benchmark for how individual receptors behave and were used to monitor activity from the manipulated changes in expression levels of paired receptors. Profiling experiments showed that the Group II mGlus exhibited faster kinetics when signaling through G_oA_ than the Group III receptors (McCullock *et al*., 2024). Monitoring the kinetics of responses (Hoare *et al*, 2020) and classical dose responses help provide a map of how responses shift as expression shifts. These parameters also allow for the detection of unique signaling properties that evolve from combined expression, especially when combined with surface expression data employed here using flow cytometry.

Our results illustrated some patterns when mGlus were stochastically assembled. For example, when mGlu2 combined with the group III receptors, the properties of mGlu2 dominated in terms of efficacy and potency, but the group III receptor kinetics were more dominant. This may be explained at least in part by the observed asymmetric roles of the subunits (Liu *et al*., 2017). mGlu7 tended to exhibit the pharmacological and kinetic properties of its dimer partner, which makes sense considering the exceedingly low potency and technically unmeasurable efficacy of the mGlu7 homodimer. This was perhaps most evident when mGlu7 combined with mGlu2 (Figs. 2 & S2), a result at least partly predicted previously (Habrian *et al*., 2019).

Some other interesting observations arose when comparing receptor expression in these experiments using flow cytometry. For example, the highest efficacy receptors, mGlu2 and 8, appeared to express at lower levels when compared to their partners. In some cases, particularly notable with mGlu7 and 8, the expression of one receptor was dominant over the other but the signaling properties favored the more weakly expressed receptor or showed intermediate properties. In general, these data indicate that at least for mGlus high efficacy does not arise from efficient expression as may be assumed, but from some other property of the receptor such as G protein coupling efficiency. These observations may help make predictions about the tendencies of each receptor to heterodimerize with specific partners compared with forming homodimers.

### Examination of mGlu heterodimers using CODA-RET2

We modified the CODA-RET system (Urizar *et al*., 2011) to examine mGlu dimers. In CODA-RET, the donor, RLuc8, is split into two fragments. Each fragment alone cannot produce luminescence, but together form a functional RLuc8 on only the desired dimer, allowing assessment of the activity of just that species through BRET with an FP tagged G⍺. Originally, this system was used to investigate D1-D2 dopamine receptors but the technology has more recently been used to isolate mGlu2/4 (Lin *et al*, 2024; Xiang *et al*., 2021) and mGlu7/8 signaling (Lin *et al*., 2022). RLuc8 is a relatively low efficiency luciferase compared to modern luciferases like NanoLuc (Hall *et al*, 2012).

Additionally, the inherent affinity between the two RLuc8 fragments is unclear. For the original studies with dopamine receptors, it was beneficial for the luciferase fragments to drive dimerization, making it easier to detect signals. This activity is not desirable for mGlus however, because mGlus drive their own stable dimerization. To circumvent this concern here, RLuc8 was replaced with Promega’s NanoBit system (Dixon *et al*, 2016), which is composed of two fragments, LgBit, an 18 kDa fragment of NanoLuc, and the SmBit peptide tag. This is essentially NanoLuc’s version of the split RLuc8, but with three key advantages. First, the activity and stability of the complemented NanoLuc was improved. Second, the NanoBit system was optimized for high light output. Finally, the SmBit peptide was optimized to have low micromolar affinity with LgBit, so there is less concern that LgBit and SmBit will drive interactions. Thus, the main driver of association would be the mGlu subunits themselves (Dixon *et al*., 2016; Schwinn *et al*, 2018). This was demonstrated in our negative control experiments using co-expression of LgBit and SmBit tags on the MOR and mGlu2, which produced only minimally detectable light compared with tagging mGlu subunits with LgBit and SmBit (Fig. S4).

The second improvement was in changing the BRET acceptor. Classically, CODA-RET is conducted using an mVenus tagged full length G⍺ protein as the acceptor, because activation of GPCRs increases the G⍺-receptor interaction. However, GPCR activation also promotes dissociation of G⍺-GTP as guanine nucleotide exchange proceeds, which means the BRET signal with CODA-RET system is an average of association and dissociation. To improve the BRET signal, mini G⍺ (mG⍺) proteins were employed. mG⍺ proteins were developed to facilitate structural studies, but have found a second life as activity probes (Wan *et al*., 2018). These proteins contain the receptor binding domains of G⍺ but lack GTP and Gβγ binding domains, so their interaction with receptors is more stable. We used an mVenus-mG⍺o construct since all mGlus exhibit high BRET signals through G_o_ proteins (Masuho *et al*, 2023; McCullock *et al*., 2024). We call this modified assay CODA-RET2 (Fig. 3).

We first used CODA-RET2 to examine orthosteric agonist responses on each combination of group II/III dimers (Fig. 3). Some surprising effects were observed. For example, mGlu2/4 showed weak potency to LY35 and L-AP4, as described previously (Kammermeier, 2012; Yin *et al*., 2014). However with mGlu2/7 and 2/8, the group II selective agonist LY35 remained a potent agonist, more potent than L-AP4 or glutamate while L-AP4 was a weaker agonist than glutamate at mGlu2/7, but not at mGlu2/8, illustrating that these three ostensibly similar dimers exhibit dramatically different pharmacologies. Further, dimers of mGlu3 and the group III receptors did not behave predictably compared with mGlu2 containing dimers.

The most dramatic and unexpected results were seen with selective allosteric modulators. While it was expected that some subtype specific compounds might be ineffective in modulating heterodimers (Kammermeier, 2012; Yin *et al*., 2014), we observed dramatically different effects of compounds on different heterodimers containing the targeted subunit. For example, the mGlu2 PAM JNJ404, which produced direct agonist effects and potentiated glutamate responses when acting on mGlu2 homodimers, produced stronger effects on when acting on the mGlu2/3 dimer. More surprisingly (Liu *et al*., 2017), when acting on mGlu2/4, JNJ404 produced weaker, but significant, direct agonist effects, but did detectably alter the activity of glutamate (see Fig. 5), suggesting that it acts as a direct agonist but not as a traditional PAM on this dimer.

When examining the effects of mGlu4 targeting PAMs, we found that they all potentiated responses of glutamate on exclusively group III containing dimers, mGlu4/4, 4/7, and 4/8, but only VU0155041 was able to modulate mGlu4/2 and mGlu4/3 in the CODA-RET2 assay (see (Fulton *et al*., 2019; Rovira *et al*., 2015)). However, the degree of potentiation on the group II containing mGlu4 heterodimers was attenuated compared to that on the purely group III containing dimers. Finally, while the mGlu7 targeting NAM ADX71743 effectively inhibited signaling of mGlu7/2 and 7/3, it only weakly inhibited mGlu7/4 and did not inhibit mGlu7/8, despite mGlu8 being the most closely related subunit to mGlu7, sharing about 80% sequence homology (Muguruza *et al*, 2016). It should be noted that the results of ADX71743 reported here differ from those reported previously (Lin *et al*., 2022). However, we examined mGlu7/8 signaling against glutamate rather than the DL-AP4 used in that study. In addition, we note that in our hands, L-AP4 produced a dramatically flattened dose-response of mGlu7/8 at concentrations ranging from 100 nM to 1 mM without reaching saturation (Fig. 3) while Lin et al. reported responses to DL-AP4 with a Hill slope near 1 that fully saturated by 10 µM, possibly indicating that there may be important differences in the CODA-RET assay and the CODA-RET2 assay used here.

Together these results highlight the complexity of mGlu dimer pharmacology and suggest that a simple binary “functional” or “nonfunctional” designation may not an appropriate characterization when considering a mGlu ligand on a particular dimer.

### mGlu homo- and heterodimers as therapeutic targets

The widespread but differential expression of the mGlus across the nervous system has made them exciting potential therapeutic targets. Their ability to modulate neuronal activity in a site or region-specific manner has long thought to be desirable neurologic conditions including epilepsy, Parkinson’s Disease, anxiety, and more. Unfortunately, this potential remains untapped, with many mGlu targeting compounds failing in clinical trials. With very few exceptions, drug screening and development campaigns hoping to leverage mGlus therapeutic potential have been homodimer centric, without consideration of the activity heterodimers. As the current study shows, this has left the mGlu field at a disadvantage. Our data demonstrating unpredictable heterodimer pharmacology, particularly with allosteric modulators, is of great concern. Without proper assessment of both homodimer and heterodimer pharmacology and physiology, the underlying mechanisms of these receptors and compounds cannot be fully understood. The present study has only scratched the surface, and although further characterization of these receptors and compounds remains a daunting hurdle to overcome, this same property offers the unique opportunity of pinpoint precision that is almost unmatched by other receptor systems. Now that the mGlu field is beginning to understand the challenges, the next steps are to systematically assess the many selective mGlu targeting compounds on each dimer species, and to identify the presence of homo- and heterodimers across the mammalian central nervous system.

## Acknowledgments

We thank Dr. Nevin Lambert (Medical College of Georgia, Augusta University) for sharing the mVenus-mG⍺o construct and for helpful discussions. We also thank Dr. Cesare Orlandi (University of Rochester) for use of his plate reader, and for helpful guidance. These studies were supported by grants R21NS126779, R03NS124987, and R01MH125849 to PJK.

