## Supplemental Info for "Unique pharmacology of mGlu homo- and heterodimers"

**Supplemental Information.**

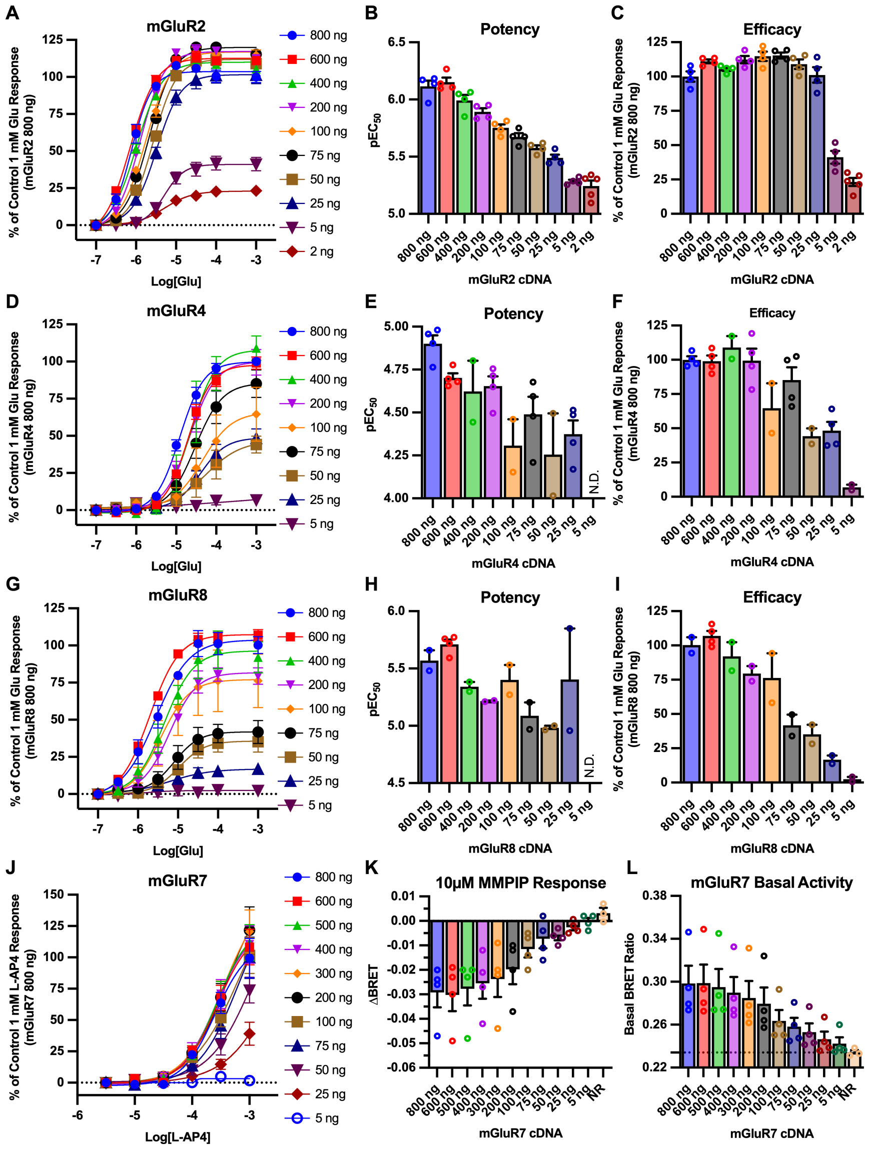


Figure S1. Effect of titration of [cDNA] levels of mGluR2, 4, 8, and 7 potency and efficacy. A-C shows glutamate does response curves (A), and summaries of glutamate potency (B) and efficacy (C) at a range of mGluR2 [cDNA], as indicated. Similar results are shown for mGluR4 (D-F), mGluR8 (G-I), and mGluR7 (J-L), but using L-AP4 as an agonist in J because glutamate responses to mGluR7 are poor. Because L-AP4 responses by mGluR7 did not reach saturation, responses to the inverse agonist MMPIP (K), which inhibits constitutive activity, and basal BRET (L) are shown as proxy measurements to estimate expression levels. Generally, decreasing mGluR [cDNA] initially reduced efficacy, consistent with a decreased receptor reserve, followed by a decrease in efficacy as [cDNA] levels were reduced further. These results are consistent with the interpretation that reduction of receptor [cDNA] coincides with reduced receptor expression levels in this assay.

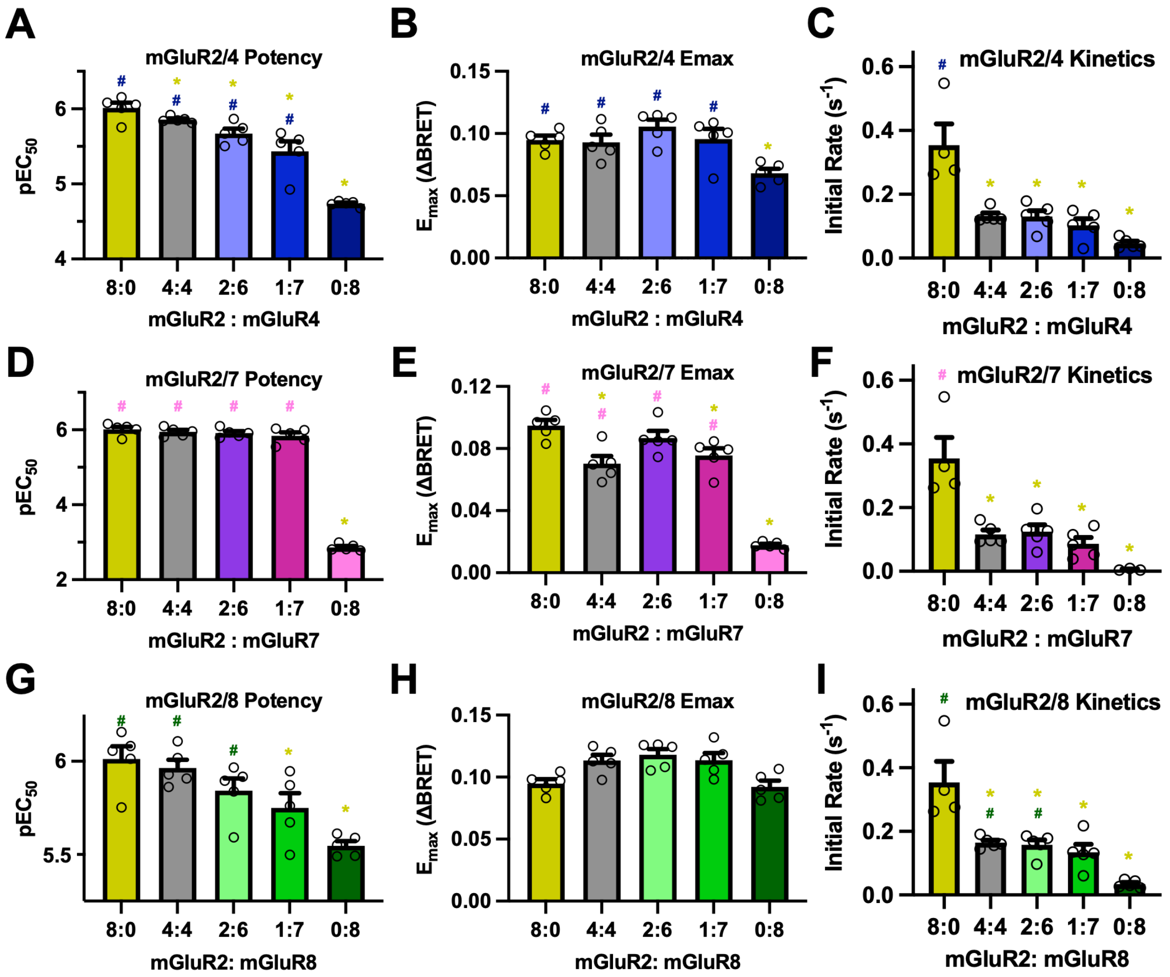


Figure S2. Summary of changes in potency, efficacy, and kinetics for stochastically assembled mGluR2-containing dimers. A-C: pEC50 values, Emax values, and initial rates of glutamate responses to hmGluR2/hmGluR4 expressed at various ratios. D-F: pEC50 values, Emax values, and initial rates of glutamate responses to hmGluR2/hmGluR7a expressed at various ratios. G-I: pEC50 values, $E_{max}$ values, and initial rates of glutamate responses to hmGluR2/hmGluR8a expressed at various ratios. Bars indicate the average ± SEM. Statistics indicate the results of a one-way ANOVA with a Holm-Šídák *post hoc* test. All conditions were compared, but for simplicity, only results of comparisons with the single expression conditions are displayed. * and # = P<0.05, when compared with the control condition of the same color as the symbol.

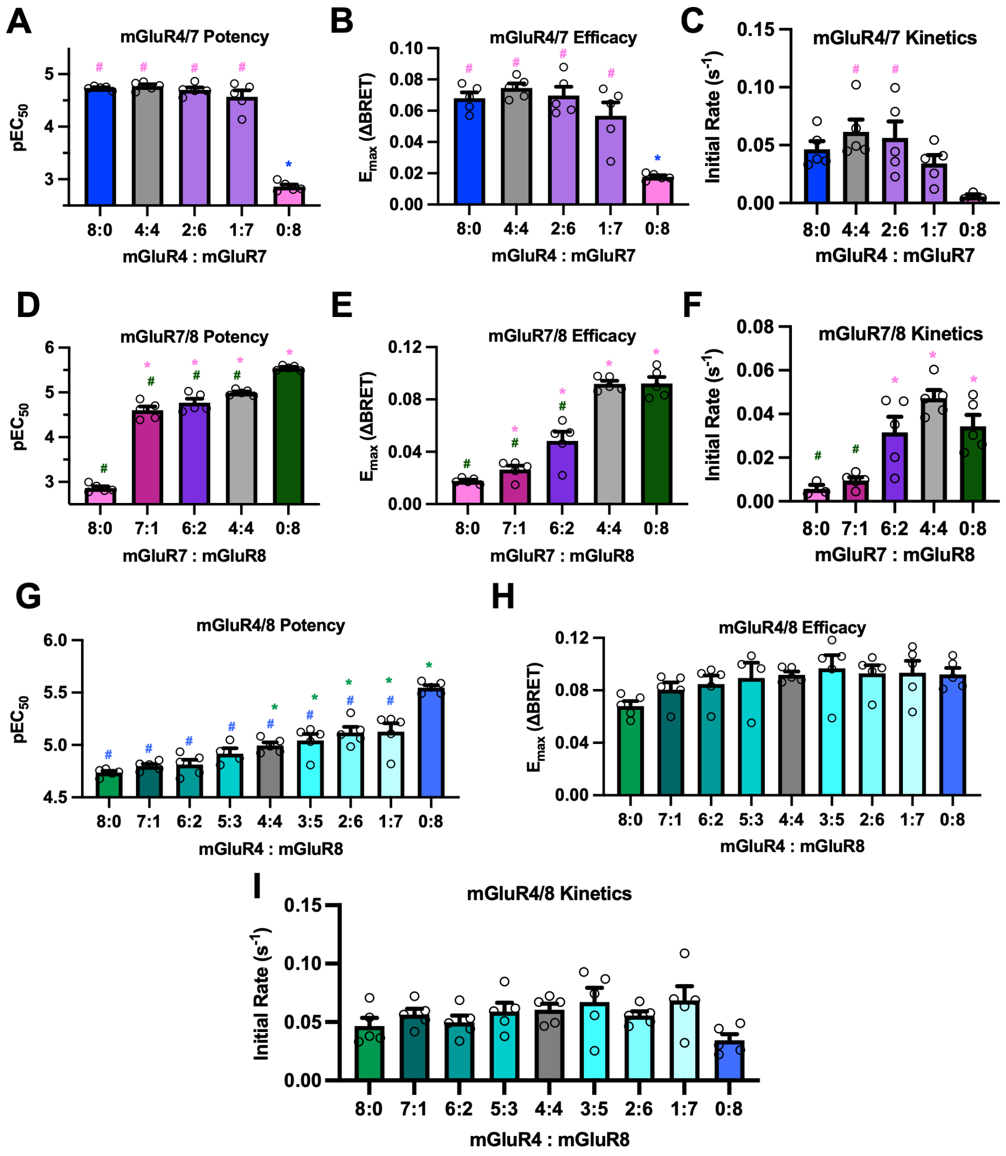


Figure S3. Summary of changes in potency, efficacy, and kinetics for stochastically assembled group III mGluR dimers. A-C: pEC50 values, Emax values, and initial rates of glutamate responses to hmGluR4/hmGluR7a expressed at various ratios. D-F: pEC50 values, Emax values, and initial rates of glutamate responses to hmGluR7a/hmGluR8a expressed at various ratios. G-I: pEC50 values, Emax values, and initial rates of glutamate responses to hmGluR4/hmGluR8a expressed at various ratios. Bars indicate the average ± SEM. Statistics indicate the results of a one-way ANOVA with a Holm- Šídák *post hoc* test. All conditions were compared, but for simplicity, only results of comparisons with the single expression conditions are displayed. * and # = P<0.05, when compared with the control condition of the same color as the symbol.

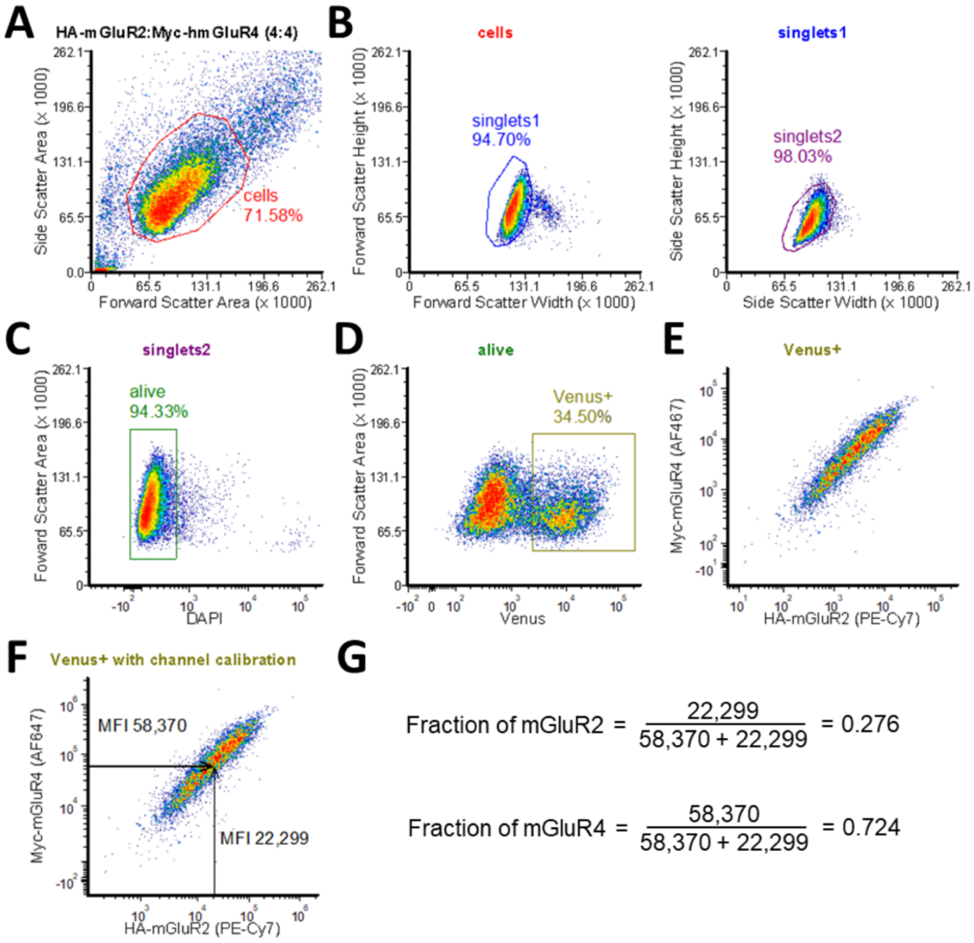


Figure S4. Flow cytometry gating scheme. A, Cells are first gated for by forward and side scatter area parameters. B, Singlets are gated for based on forward and side scatter height and width parameters. C, Live cells are gated based on their ability to exclude DAPI. D, Transfected cells are gated based on Venus fluorescence from the co-transfected split-Venus on Gbg sububits. E, Surface expression of HA-mGluR and myc-mGluR (2 and 4, respectively in this example) is detected using fluorescently conjugated primary antibodies against each epitope. F, Surface expression after channel calibration with Quantum Simply Cellular beads. G, Median fluorescence intensity values (MFI) for the two channels are used to determine the fraction of each receptor on the cell surface.

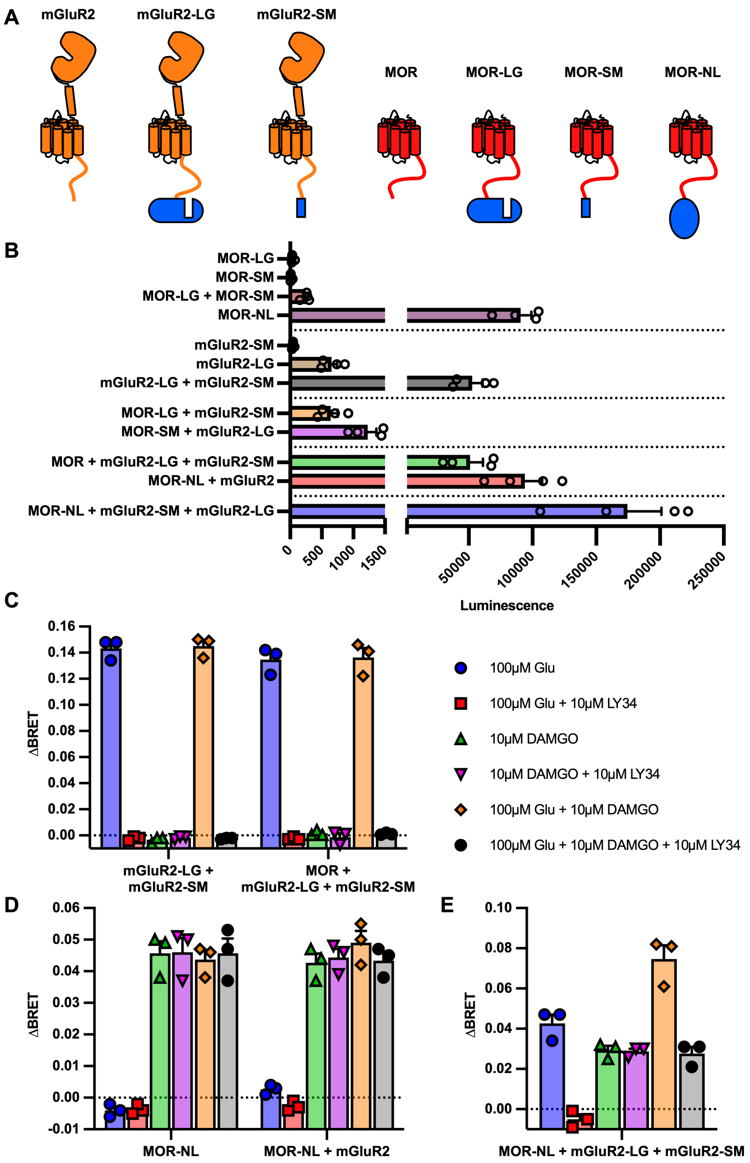


Figure S5. Control responses in the CODA-RET2 system. A, summary of the constructs used for CODA-RET2 control experiments. B, Luminescence responses in the presence of the indicated receptor constructs. Note the split x-axis. While a small amount of luminescence was detectable when only LgBit constructs were expressed, robust expression was only observed when mGluR2-LgBit and mGluR2-SmBit were coexpressed, or when a full length NanoLuc was expressed. C-E, Summary of delta BRET values upon application of the indicated agonist(s). When mGluR2-LgBit and mGluR2-SmBit were coexpressed, only glutamate resulted in a detectable delta BRET, even when MOR was coexpressed (C), and this response was blocked by the mGluR2 antagonist LY34. With MOR-NLuc expression, only DAMGO resulted in a detectable delta BRET (D). Finally, with coexpression of MOR-NLuc, mGluR2-LgBit, and mGluR2-SmBit, cells responded to both glutamate and DAMGO, and the glutamate responses were blocked by LY34, while the DAMGO responses (or portions of responses to combined agonists) were not (E).

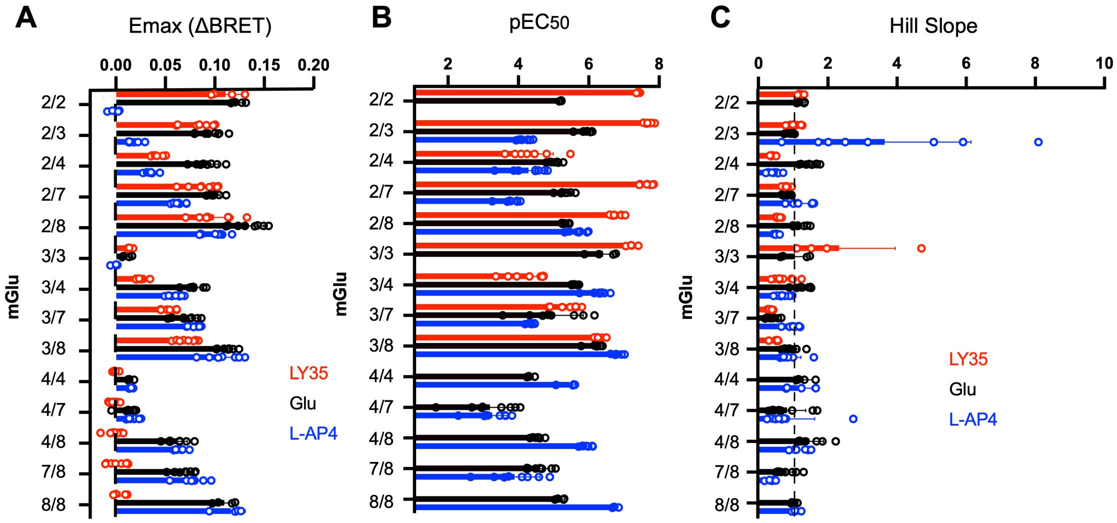


Fig. S6. Summaries of Emax values (A), pEC50 (B), and Hill slopes (C) from the dose-response data shown in Fig. 3. Bars represent average ± SEM and individual measurements (open circles) for each of the indicated mGlu heterodimer. No symbols are shown for values that could not be calculated due to lack of responses.

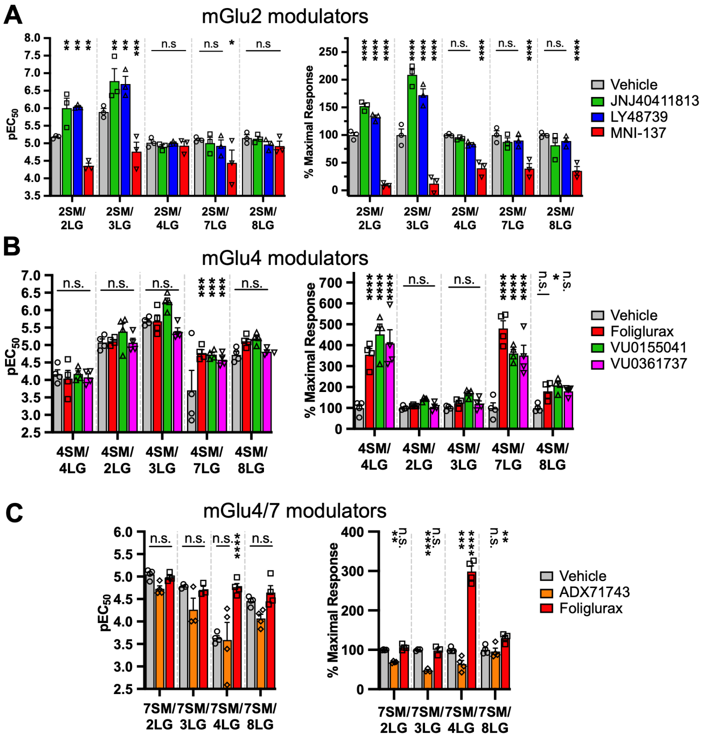


Figure S7. Panel A shows average ± SEM $pEC_{50}$ (left) and Emax (right) values for the experiments shown in Fig. 4A. Panel B shows average ± SEM $pEC_{50}$ (left) and Emax (right) values for the experiments shown in Fig. 5A. Panel C shows average ± SEM $pEC_{50}$ (left) and Emax (right) values for the experiments shown in Fig. 5D.
